# A critical review of the use and performance of different function types for modeling temperature-dependent development of arthropod larvae

**DOI:** 10.1101/076182

**Authors:** Brady K. Quinn

## Abstract

Temperature-dependent development influences production rates of arthropods, including crustaceans important to fisheries and agricultural pests. Numerous candidate equation types (development functions) exist to describe the effect of temperature on development time, yet most studies use only a single type of equation and there is no consensus as to which, if any model predicts development rates better than the others, nor what the consequences of selecting a potentially incorrect model equation are on predicted development times. In this study, a literature search was performed of studies fitting development functions to development of arthropod larvae (99 species). The published data of most (79) of these species were then fit with 33 commonly-used development functions. Overall performance of each function type and consequences of using a function other than the best one to model data were assessed. Performance was also related to taxonomy and the range of temperatures examined. The majority (91.1 %) of studies were found to not use the best function out of those tested. Using the incorrect model lead to significantly less accurate (e.g., mean difference ± SE 85.9 ± 27.4 %, range: −1.7 to 1725.5 %) predictions of development times than the best function. Overall, more complex functions performed poorly relative to simpler ones. However, performance of some complex functions improved when wide temperature ranges were tested, which tended to be confined to studies of insects or arachnids compared with those of crustaceans. Results indicate the biological significance of choosing the best-fitting model to describe temperature-dependent development time data.

**Highlights:**

- Temperature-dependent development functions of arthropod larvae were reviewed
- 79 published datasets were re-tested and fit with 33 different function types
- 91.1 % of published studies did not fit their data with the best function of those tested
- Performance differed among functions and was related to taxon and temperature range tested
- Function type impacted predicted development times, so using the best function matters

## 1. Introduction

Temperature affects biota at all levels, ranging from effects at the fundamental biochemical and physiological levels (Bĕlehrádek, 1935; Coutant and Talmage 1976; Somero, 2004) to effects on individual organisms (Brière et al., 1999; MacKenzie, 1988), populations (Aiken and Waddy, 1986; Cooper et al., 2012; McLaren et al., 1969), communities, and ecosystems (Menge, 1978; McQuaid and Branch, 1985). Through its effects on the physical and chemical properties of biologically active molecules, such as enzymes, temperature affects the rate at which numerous life processes occur, including metabolism, oxygen consumption, photosynthesis, movement, survival, growth, and embryonic development (Bĕlehrádek, 1935; Brière et al. 1999; Corkett, 1972; Coutant and Talmage 1976; Du et al., 2007; Geffen and Nash, 2012; Herzig, 1983; McLaren et al., 1969). Temperature also significantly affects larval development rate of poikilothermic animals, including some vertebrate larvae (Lind and Johansson, 2007; Kang et al., 2009; Miller et al., 2006) and those of invertebrates (e.g., de Severyn et al., 2000; Jenkins et al. 2006; Singh and Sharma, 1994). Temperature has particularly strong impacts on moulting and development of arthropods (Anger, 1984; Corkett and McLaren, 1970; Easterbrook et al., 2003; Hamasaki et al., 2009; Koda and Nakamura, 2010; MacKenzie, 1988; Marchioro and Forester, 2011; McLaren, 1963).

Within certain tolerance limits (Bĕlehrádek, 1935; Brière et al., 1999; Campbell et al., 1974; Shi and Ge, 2010) rates of biological processes of poikilotherms, including larval development, are positively correlated with temperature; thus, higher temperatures generally result in more rapid development than lower temperatures. This has important ecological implications, as environmental temperatures can influence generation times, production cycles, and population dynamics of such organisms. Higher or lower temperatures could, for example, lead to changes in the amount and/or timing of peak secondary marine production of copepods (Huntley and López, 1992; McLaren, 1963), outbreaks of agricultural pests (Easterbrook et al., 2003) or vector-borne diseases (Bayoh and Lindsay, 2003), or introduction and establishment of invasive species into new areas (de Rivera et al., 2007). Water temperatures could also influence patterns of recruitment to populations of marine invertebrates, including crustaceans such as lobsters and crabs, on which human fisheries depend (Aiken and Waddy, 1986; Anger, 1984; Caddy, 1986; MacKenzie, 1988; Rothlisberg, 1979).

When modeling ecology and population dynamics of arthropods, equations are used to represent the functional relationship between environmental temperature and development rate or time of larvae. These equations, hereafter referred to as development functions, are derived by rearing larvae at different controlled temperatures in a lab or hatchery setting, observing development times of multiple larvae at each temperature, and then using regression analyses to fit an equation relating temperature to development time (or its inverse, rate) to the data obtained. There are countless potential forms of equation that can be used to fit such data, including various linear, simple curvilinear, and complex non-linear functions (e.g., see reviews by Anger, 2001; Angilleta Jr., 2006; Blanco et al. 1995; Guerrero et al., 1994; Heip, 1974; Kontodimas et al., 2004; McLaren, 1995; Shi and Ge, 2010; Smits et al. 2003). These functions differ in form, assumptions, procedures used to derive their parameters, and most importantly in terms of the development times predicted. For example, development times of American lobster, *Homarus americanus* (H. Milne Edwards, 1837), larvae predicted with 33 of these development function types can differ from each other and the data used to derive them by ≥ 50 days at the same temperatures (see Fig. 1; Table S1). However, this is no clear standard rule or consensus as to what is the “best” type of development function to apply to these kinds of data. Researchers are generally left to choose the type of development function to use on their own, and will often select one or a very few forms that have the best apparent match to their characters of their data or has been used by other studies on the same or related species (e.g., Edgar and Andrew, 1990; McLaren et al., 1969). Given the potential for different functions to make very different predictions of development times (e.g., Fig. 1), however, development function choice should be given more consideration in studies on these species.

**Figure 1.**
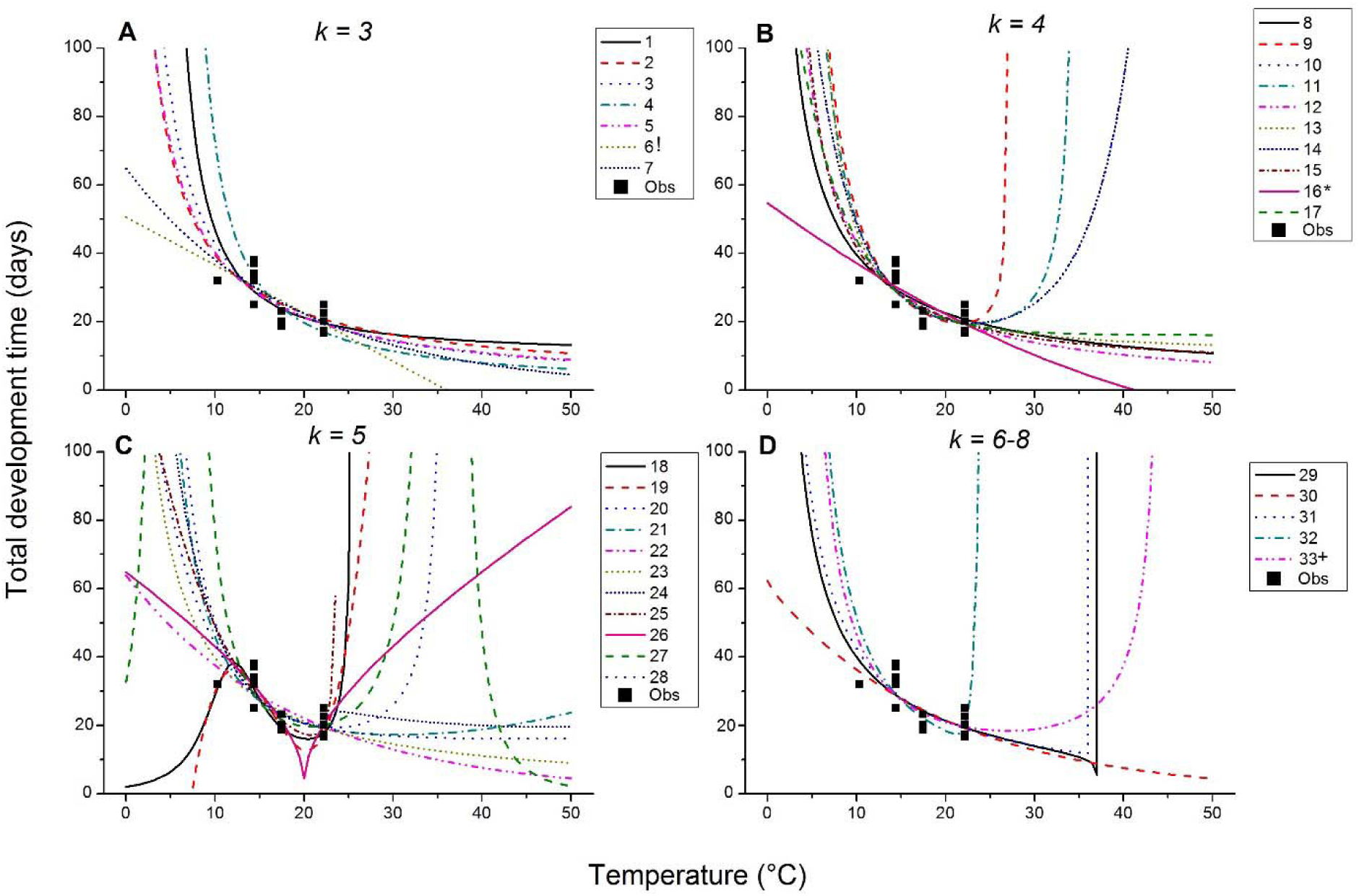
Examples of the curves formed by different development functions, when these are used to calculate development time (y-axes, days) at different temperatures (x-axes, °C). Types of functions plotted here are those presented in Table 1, separated into those with *k*-values (number of parameters + 1) of (A) 3, (B) 4, (C) 5, or (D) 6-8. Actual functions (see Table S1) were derived from and fitted to data for total development times (combined time to complete larval stages I-III) of American lobster (*Homarus americanus* (H. Milne Edwards, 1837)) larvae, as reported by Quinn et al. (2013); observed development times are plotted (squares) along with predictions of development functions (lines). Coefficients, R^2^, AIC_C_, and Δi values for these functions are presented in Table S1. ‘*’ = function used by Quinn et al. (2013) to fit the data, ‘!’ = actual “best” function for these data, and ‘+’ = worst function for these data.

It is possible that certain function types may in general be better representations of the relationship between temperature and development of arthropod larvae, or specific sub-groups within the Arthropoda (e.g., arachnids vs. crustaceans vs. insects), for example because they come closer to capturing thermal performance relations of enzymes and other biomolecules mediating moulting and development cycles in these taxa (Brière et al., 1999; Huey and Stevenson, 1979; Somero, 2004). As a result, such functions might also achieve better fit to development data, be able to more-closely match real observed development times, and make better predictions of development in nature. Differences in methodologies used in studies on different taxa, for example the fact that the range of temperatures tested is generally wider for insects and arachnids than crustaceans; (reviewed by Hartnoll, 1982; Quinn and Rochette, 2015), might also lead to apparent taxonomic differences in function performance and should be investigated. Several previous studies have compared the characteristics of different development function types in general (Anger, 2001; Blanco et al., 1995; Guerrero et al., 1994; McLaren, 1995). Others have examined performance of one or two specific functions on multiple species (e.g., Logan et al., 1976), or attempted to fit multiple function types to data for one or two specific species under study to select the best function for their data (e.g., Angilletta Jr., 2006; Heip, 1974; Kontodimas et al., 2004; Shi and Ge, 2010; Smits et al., 2003). Many other studies seem to choose one or very few function(s) semi-arbitrarily, without discussing alternatives (e.g., de Oliveria et al., 2009; Thompson 1982; see also Results). However, no previous study has attempted to assess the degree to which one versus multiple types of development functions are used in published studies, compared performance of different types of development functions across multiple species, or assessed the overall impact of function choice to predictions made with such functions. Such a large-scale analysis is needed, though, because it could potentially allow functions that tend to better represent development data in general to be identified, which can then allow for more informed decisions by future studies on arthropod larval development.

In this study, a critical literature review was conducted to assess whether and to what extent studies of temperature-dependent development of arthropod larvae attempt to represent their data with more than one development function type, and also which specific types of functions tend to be used. Then, data from previously published studies were extracted and retested to derive multiple different development functions for the same datasets. The best model type for each dataset was determined, and whether or not published studies actually used the best function type for their data was recorded. Overall performance of different function types were assessed by comparing overall function rankings, proportion of variance explained, and information loss across datasets. Any taxonomic patterns (e.g., whether particular function types performed better for arachnids than for crustaceans and/or insects) and whether performance was related to the range of temperatures tested were also noted. The consequences of using different function types were then tested by comparing the difference of predicted versus observed development times, fit, and information loss of the function type used in original studies versus that of the best function for the data.

## 2. Materials and methods

### 2.1 Literature review

A literature search was conducted through Web of Science (Thompson Reuters, 2015) for the terms “temperature” AND “development”. An initial search was carried out on 12 September 2012, through which the majority of the data in the present study were obtained; this search yielded 1,052 results. A second search was carried out on 19 November 2014, which returned 35 additional results not available or published online at the time of the initial search. These 1,087 total search results were them further examined, and several criteria were used to remove non-relevant results. Accessible peer-reviewed studies that reported larval development rates or times of arthropods at different temperatures and derived a regression equation(s) (i.e., development function) from their data were sought out. Studies that looked exclusively at growth (size increase), which is a distinct process from development (Forster et al., 2011), were excluded.

After applying these criteria, 81 studies of 96 different arthropod species were obtained, which provided a total of 99 species datasets for subsequent examination and analyses (Table S2). Several specific types of development function were frequently utilized in these studies (Table S2); these functions are presented in Table 1 and discussed in the next section (2.2.1, below). Studies were published between 1970 and 2014, and conducted in several different countries on marine, freshwater, and terrestrial species (Table S2) of various taxa within the Arachnida (Phylum Arthropoda: Subphylum Chelicerata), Crustacea, and Insecta (Table S2). To assess whether published studies tested multiple development functions on their data, each study was carefully read and the number and types of development functions used to fit the data for each study species were noted. Even if results from multiple functions were not reported, if alternative functions than reported were at least mentioned in the Methods sections of studies they were counted as having considered > 1 function. Also if any study tested multiple development functions on their data and concluded one of these to be the “best” function for their data this was also noted. The percent (%) of the 99 species datasets from the literature search on which one, two, or more functions were tested, and the % of datasets on which different types of functions were tested, were then calculated.

**Table 1.**
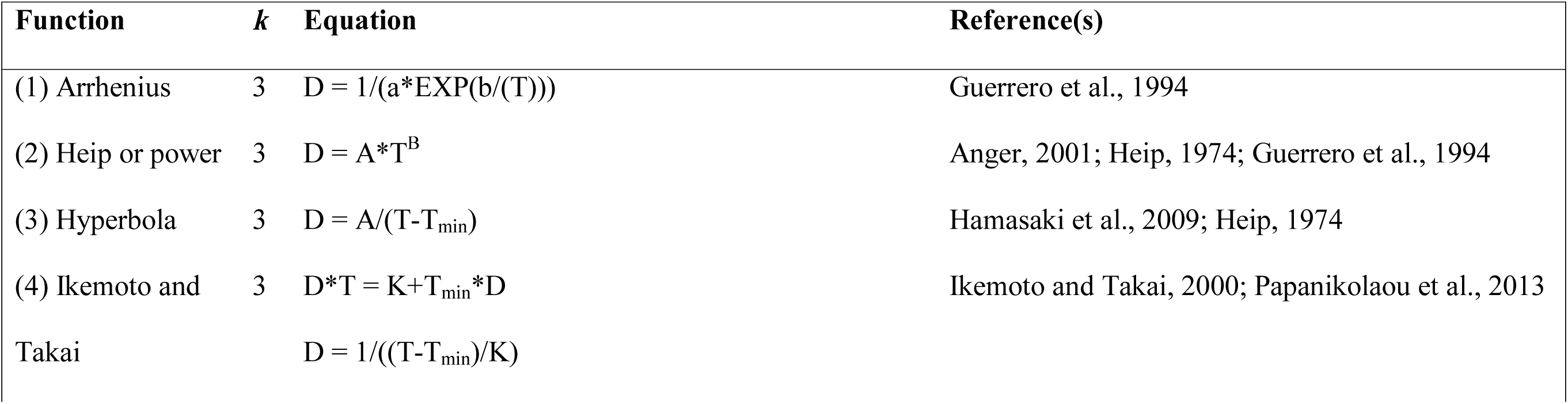
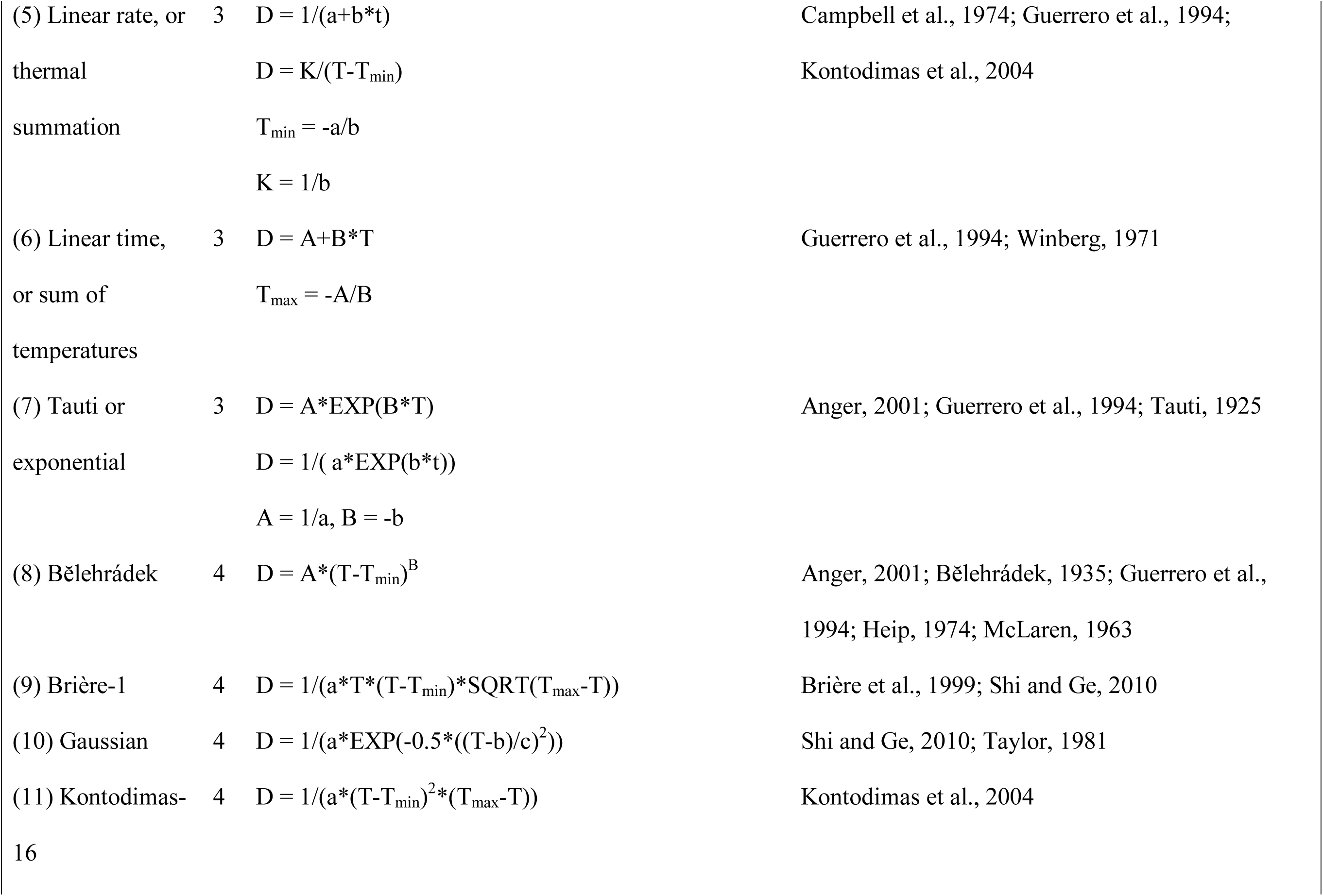
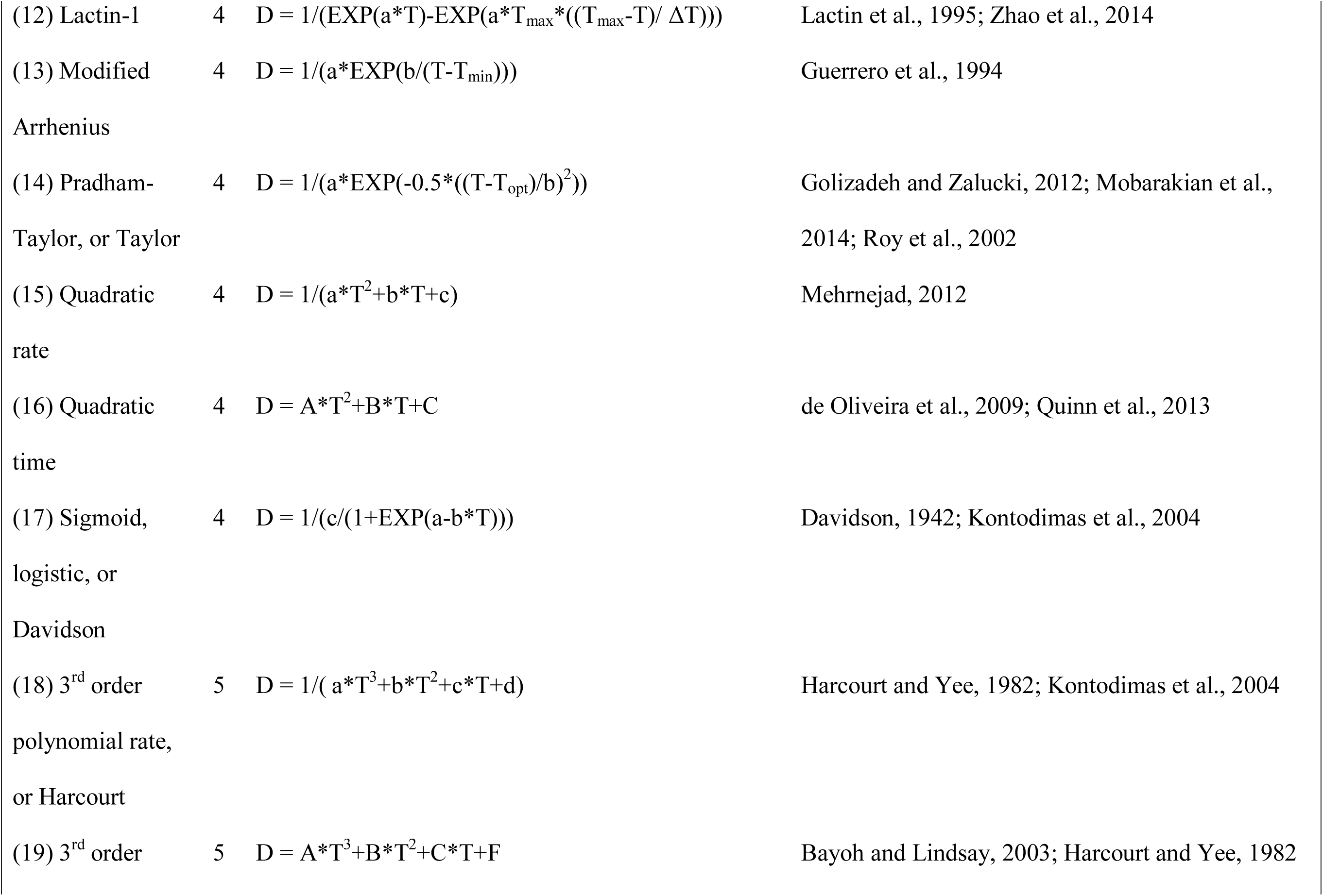
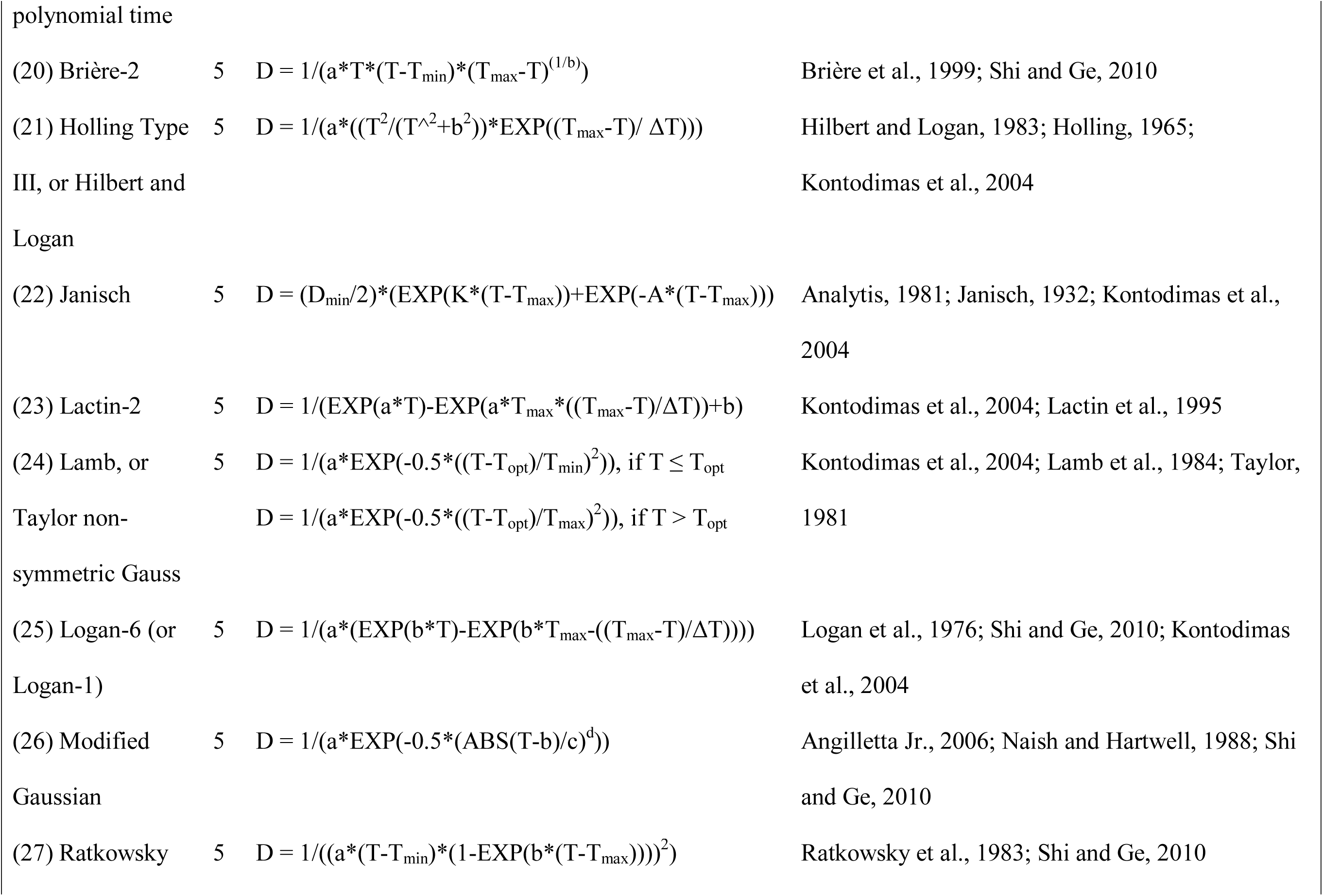
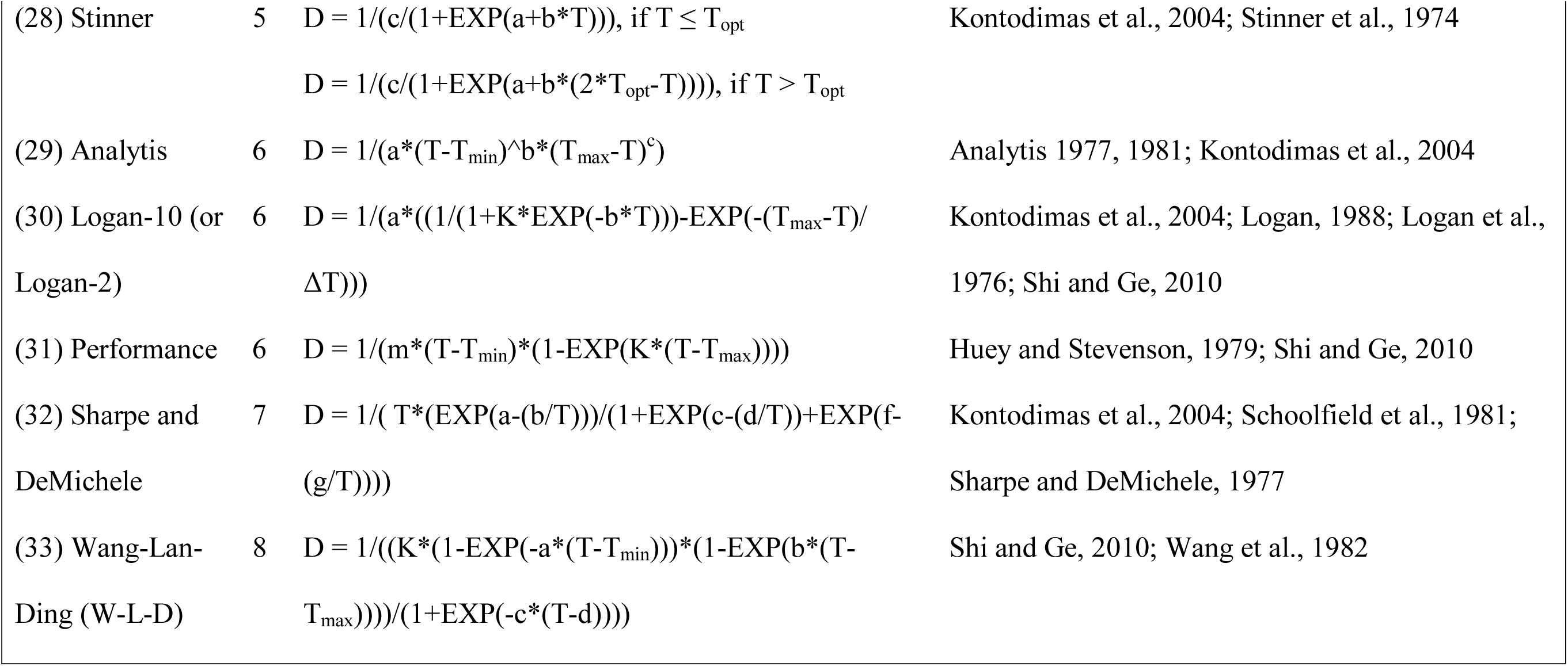
Names and forms of the 33 different development functions examined in this study. Functions are arranged and numbered first in order of their value of *k* (number of parameters + 1, used in AIC_C_ analyses), and then alphabetically by their common name. In all equations D is development time (in days), T is temperature (°C), A, B, C, F, a, b, c, d, f, and g are fitted constants. Additional fitted constants in some equations with biological meaning and constrained values are as follows: T_min_ = biological minimum temperature; T_max_ = biological maximum temperature; T_opt_ = temperature at which development rate is maximized; K = thermal constant, or the number of degree days required to complete development (Kontodimas et al., 2004); ΔT = range of temperatures over which development occurs; D_min_ = minimum development time. Functions that are fitted to development rate (1/D) are here represented in their inverted form, from which D can be directly calculated. For clarity lowercase letters are used for functions fitted to development rate, and uppercase letters are used for functions fitted directly to time (D).

### 2.2 Meta-analysis

#### 2.2.1 Development functions considered in this study

In the present study, 33 development function types were examined (Table 1; Figure 1). These functions were used because they were found in the literature review in the present study (see section 2.1 and Results) to be used quite frequently in studies of arthropod larvae, and were discussed in reviews and studies on the topic of arthropod temperature-development functions by Anger (2001), Guerrero et al. (1994), Heip (1974), Kontodimas et al. (2004), and Shi and Ge (2010). Three linear and 30 nonlinear functions were examined, with *k*-values (*k* = numbers of parameters + 1; Anderson, 2008) ranging from 3 to 8 (Table 1). Eight of these functions are fit directly to development time data, while the remaining 25 functions are typically fitted to development rate data, which are the inverse of time (Table 1); in some cases, the same function form (e.g., quadratic) is applied to either development rate (function #15, Table 1) or time (#16), yielding distinct functions (Fig. 1). 12 of these functions included a nonzero minimum temperature (T_min_) for development at which development rate is zero and development time becomes infinite (see Fig. 1), 15 included a similar maximum temperature for development (T_max_), and 8 functions included both of these thresholds (Table 1). Linear functions can be used to derive starting estimates for the values of T_min_ and T_max_ (Campbell et al., 1974; Kontodimas et al., 2004; Table 1) to be used in deriving more complicated nonlinear functions. Attempts were also made to test three additional functions found in these reviews: the Exponentially Modified Gaussian (Naish and Hartwell, 1988), Sharpe-Schoolfield-Ikemoto (SSI; Sharpe and DeMichele, 1977; Schoolfield et al., 1981), and Weibull (Angilletta Jr., 2006) functions. However, these three functions have very complex structures requiring specialized fitting procedures not readily applicable in many statistical software packages (see review by Shi and Ge, 2010), and in the present study they could not be fit to the datasets used; as such, they were not considered further in this study.

#### 2.2.2 Re-analysis of species datasets from published studies

To assess whether the function(s) used in published studies were actually the “best” functions for these published datasets, data were extracted for re-analysis from as many of the studies obtained through the literature search described above as possible. Studies from the literature search that did not present their data in a way that allowed it to be extracted for retesting (e.g., only mean development times resented, without any measure of error), had to be excluded. Therefore, only 79 species datasets out of the 99 initially obtained from the literature search could be retested (Table S2); these included 10 arachnids (9 mites in the Subclass Acari and one spider in Subclass Aranea), 28 Crustaceans (9 copepods and 19 decapods), and 41 insects in 8 orders (8 Coleoptera, 6 Diptera, 9 Hemiptera, 1 Homoptera, 1 Heteroptera, 8 Hymenoptera, 6 Lepidoptera, and 2 Thysanoptera) (Table S2). Raw data or means ± error (standard deviation (SD), standard error (SEM), etc.) and sample sizes were extracted from tables or figures in published papers for each of these 79 species and used to generate datasets for reanalysis.

Each study dataset was analyzed with linear and nonlinear regressions (Table 1) between temperature and development time or rate, as appropriate. These regression were carried out using IBM SPSS Statistics 22 (SPSS Inc., 2014). To simplify analyses, only total development times or rates (i.e., summed across multiple larval stages) were examined and data for individual stages were not. For development functions including thermal limits or other parameters with biological meaning (e.g., T_min_ or T_max_; see Table 1 and section 2.2.1), unconstrained regressions were initially carried out, with starting values for these parameters set to values estimated from linear functions (see Table 1 and section 2.2.1, above). However, results were not accepted if this yielded biologically unrealistic estimates, such as T_min_ < 0°C in a species not known to survive and develop at sub-zero temperatures, unreasonably low T_min_ (e.g., −100°C) or high T_max_ (e.g., 100°C), or T_min_ or T_max_ within the temperature range for which successful development was reported in the original study. In this case, constrained regressions were carried out (e.g., T_min_ ≥ 0°C, T_min_ < minimum T with successful development, T_max_ > maximum T with successful development, etc.) until satisfactory values were obtained.

Once a regression equation corresponding to each development function was obtained for each dataset, AIC_C_ values (Akaike’s Information Criterion (AIC) corrected for finite sample size; Akaike, 1973; Anderson, 2008) were calculated using the residual sum of squares (RSS) between observed development times and those predicted by each function for each study (Anderson, 2008). Development functions were then ranked for each species dataset based on AIC_C_-values. The “best” possible rank was 1, corresponding to the lowest AIC_C_ value, and the “worst” possible rank – if there were no ties – was 33, which corresponded to the highest AIC_C_ value among the functions tested. The percentage of retested datasets for which each of the 33 functions was concluded to be “best” was recorded. For each species dataset, whether or not the function used in its original published study and concluded to be the “best” function for the data was the same as that determined to be the “best” function in this study was noted. If more than one function was used in an original study, whether the actual best function was among all these functions was also noted. RSS values were also used to calculate R^2^-values for each function, on each dataset. These values were used as a measure of function performance (see next section, 2.2.3), indicating the proportion of the variation in observed development times that was explained by each temperature-dependent development function.

#### 2.2.3 Assessing and comparing the overall performance of different functions

To determine whether any particular type(s) of function tended to do “better” than others, the overall performance of each development function across species datasets was assessed using three measures: average ranking (determined using AIC_C_), R^2^, and Δ_i_ values. These measures of performance were also compared across different taxonomic groups (arachnids, crustaceans, and insects) as it was possible that certain functions might better represent development of animals in a certain group(s) better than animals in others; this could be due to real biological differences among taxa or to different experimental methodologies used for their rearing.

Calculation of function rankings and R^2^-values for each function on each dataset was described above (previous section). Δ_i_ values were calculated once the best function for a given dataset was determined to assess the information potentially lost by using a function other than the best one (Anderson, 2008); a lower Δ_i_ value is better, and indicates less information loss. The Δ_i_ value for a given function, “i", is calculated by subtracting the AIC_C_ value of the best function of those tested from its AIC_C_ value (Anderson 2008); therefore, the best function will have Δ_i_ = 0. A function with a lower Δ_i_, higher R^2^, or lower (better) ranking value on average than other functions was considered to have performed better overall than other functions.

Separate two-way ANOVAs were carried out in IBM SPSS Statistics 22 (SPSS Inc., 2014) to compare each of these three measures of performance (rankings, R^2^, and Δ_i_) across different development functions (factor with 33 levels; Table 1), as well as among different taxonomic groups (factor with 3 levels: Arachnida, Crustacea, and Insecta). R^2^-values were arcsine-square root transformed to meet the assumptions of parametric tests. If a statistically-significant (p ≤ 0.05) interaction between function and taxon was found, the data for that measure were split by taxon and then separate one-way ANOVAs comparing different functions were carried out for each taxon. If significant differences among functions were found, Tukey’s Honestly Significant Difference (HSD) test was used to perform post-hoc comparisons among specific function types.

#### 2.2.4 Assessing the consequences of development function choice

The consequences of choosing one function versus another to predict larval development at different temperatures were assessed by comparing whether and how much the best or only function originally used in the study from which each species dataset was obtained (“original best” function) predicted observed development times relative to the best function identified in this study (“actual best” function). Three measures, described in detail below, were calculated for each of the 79 species datasets to assess consequences of function choice. These were mean error, R^2^ decrease, and Δ_i_ resulting from using the original study function instead of the actual best one.

To calculate the first of these, error, the absolute deviance (in days) between predicted (using a development function) and observed development time was calculated for both the best and original study functions, at each temperature tested in original studies. The absolute deviance for the actual best function was then subtracted from that for the original best/used function at each temperature, to determine how much predictions were worse (i.e., further from observed values) when using the original versus best function. These differences were then averaged across all temperatures and data points to calculate a mean absolute error (in days) per each dataset that was due to using the original versus actual best functions. Mean error per dataset was also expressed as a percent improvement by performing the aforementioned calculations, but before averaging differences between deviance of original and actual best functions across temperatures these differences were divided by the best function’s deviance and multiplied by 100 %; this translated the error from a “raw” measure (in days) to a percentage (%).

R^2^ decrease was simply calculated for each dataset by subtracting the R^2^-value of that dataset’s actual best function from that of its original used function. Percent R^2^ increase was also calculated for each dataset by dividing the R^2^ change by the best function’s R^2^, and then multiplying by 100 %. A large R^2^ decrease implied that a lower proportion of the variation in development time was explained by the original function than the best one.

The Δ_i_ of the original function for each dataset was also examined to assess the extent of potential information lost by using these, rather than the best functions, to fit the data. As described above, a lower Δ_i_ (closer to 0 = Δ_i_ of best function) is better, and implies the function retains more useful information than a function with a higher Δ_i_. Generally a function with Δ_i_ < 2 contains some useful information, even if it is not the “best” function, whereas a function with Δ_i_ > 14 is highly unlikely to be informative (Anderson 2008). A high original function Δ_i_ value would imply that the original function was considerably less informative than the best function.

If the best function for a given dataset was the same as the original function, all of the measures described above would have a value of zero; if any measure were not significantly different from zero, then, it would imply that using the best versus original function did not result in meaningfully different predicted development times. Therefore, the distributions of each measure of the consequences of using a function other than the best one (mean errors and R^2^ decrease (both raw and % versions), as well as original function Δ_i_ values) across all species datasets were compared to zero (null hypothesis of no differences) using five one-sample *t*-tests. If the null hypothesis comparing these data against zero-values could be rejected, then a significant (p < 0.05) impact of function choice was concluded.

### 2.3 Potential relationship between thermal range and function performance

One interesting pattern noted during the literature review in this study was that studies of temperature-dependent development differed considerably in their methodology, particularly regarding the range of temperatures tested. Thermal ranges varied considerably among studies in general, from as narrow as 6°C to as wide as 38°C (Table S2). Differences also appeared to exist between studies of different taxonomic groups, particularly between studies of crustaceans versus those of arachnids and insects (Hartnoll, 1982; Quinn and Rochette, 2015; Table S2). To confirm whether such taxonomic differences were significant, the thermal range for each species dataset obtained in the initial literature search (n = 99 total) was calculated as the maximum temperature tested in its original study minus the minimum temperature tested; this included any temperatures at which successful development was not observed (i.e., survival = 0 %), as this implies a developmental threshold (i.e., T_min_ and T_max_; see Table 1). One-way ANOVA was then used to compare thermal ranges among the three major taxonomic groups of arthropods.

More complex functions with more parameters have lower power to model smaller datasets (Angilletta Jr., 2006; Shi and Ge, 2010). This is especially true for functions containing threshold temperatures (T_min_ and T_max_) if these are fit to data recorded over narrow thermal ranges not approaching a species’ real thermal limits (Shi and Ge, 2010). In such cases, thermal thresholds have to be extrapolated too far beyond actual observations, resulting in excessively extreme estimates for these parameters. One could thus expect that more complex functions may perform better when wider thermal ranges are tested, potentially approaching or encompassing real thermal limits. To test whether or not more complex functions would be selected as better functions when wider thermal ranges were tested, Pearson’s correlation coefficients (R) were calculated between the thermal ranges calculated for each dataset and the ranking of each function per dataset. Thus, 33 separate correlation analyses were carried out (one for each type of development function), each consisting of 99 thermal range-ranking pairs (one pair per dataset). Whether correlations were significant (p ≤ 0.05), and if significant R-coefficients were positive or negative was examined. As lower ranking values implied better function performance (see above), a positive correlation for a given function implied that the function did worse when a wider range of temperatures was tested, whereas a negative correlation meant that the function did better when a wider thermal range was tested.

## 3. Results

### 3.1 Results of literature review – usage of different functions

Out of 99 different species datasets, over half (59.6 %) were reported to have been fit with only one development function and the vast majority (96.0 %) were fit with five or fewer functions (Fig. 2). In most cases studies that used 2-5 functions examined insects or arachnids (Fig. 2) and used 1-3 more complex functions plus the linear rate (function #5) or Ikemoto and Takai (#4) functions to derive starting values for T_min_ and T_max_ parameters in these complex functions (Table S2; see also below and Fig. 3). The only taxonomic group for which > 5 different functions were tested was Insecta, for which 4.0 % of all species datasets (representing 7.8 % of insects) were tested with 5-17 different development functions (Fig. 2).

**Figure 2.**
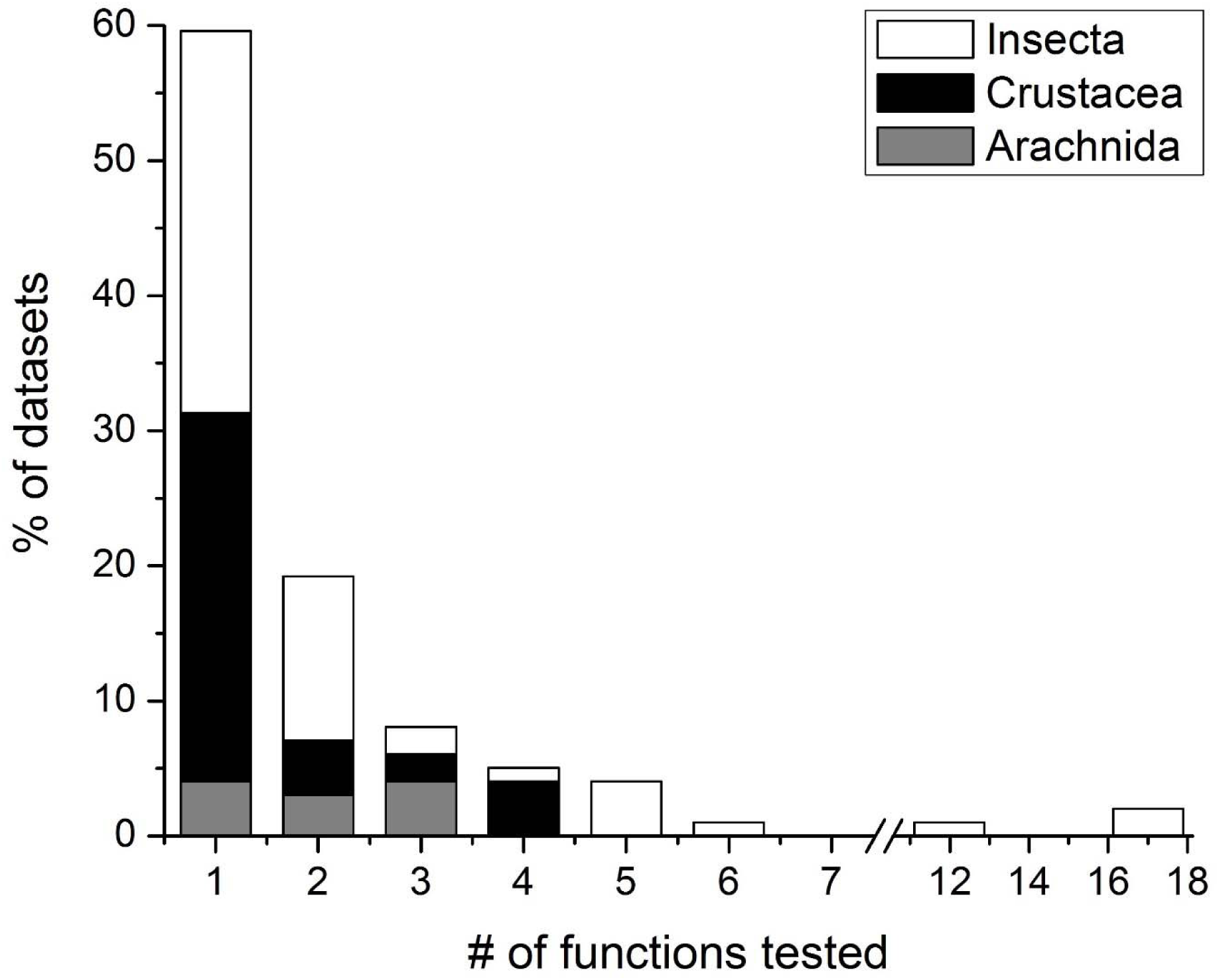
Usage in each development study reviewed of one or more different types of development function. The percentage (%) of species datasets obtained in initial literature review (n = 99 total, see Table S2) that were tested with each number of functions is plotted on the y-axes and broken down by taxa (gray bars = arachnids, black = crustaceans, and white = insects). For names and details of development functions (#1-33) the reader is referred to Table 1.

**Figure 3.**
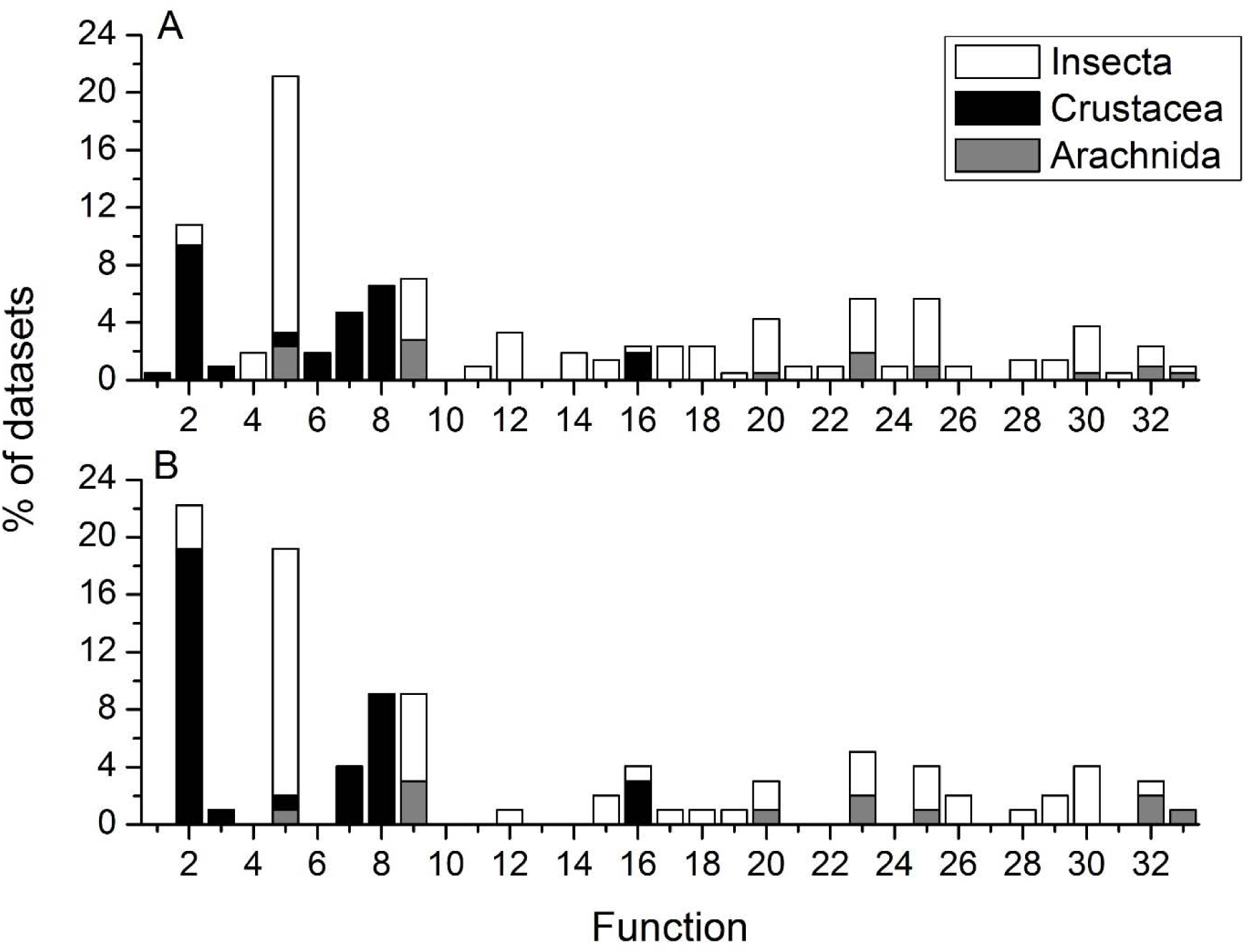
Usage in each development study reviewed of specific development function types. The percentage (%) of the 99 species datasets obtained in initial literature review (see Table S2) that were (A) tested with each type of function (i.e., all used functions) and (B) concluded to be best represented by each type of function (i.e., best used function) is plotted on the y-axes.Results are also broken down by taxonomic group, as in Fig. 2. For names and details of development functions (#1-33) the reader is referred to Table 1.

The most frequently-used development function overall, and particularly among studies of the Insecta and Arachnida, was the Linear rate function (#5), which was used in 34.9 % of all datasets and representing 28.4 and 15.2 % of insect and arachnid datasets, respectively (Fig. 3A; Table S2). Seven other function types were used for arachnids, and for insects a wide range of 24 further functions types were used (Fig. 3A). Functions used on insects and arachnids ranged in complexity from relatively simple (*k* = 3) to very complex (#33, *k* = 8), but with no particular function aside from the linear rate one (#5) predominating (Fig. 3A). Studies of Crustacea used a more limited set of 8 functions, all but one of which (function #16, *k* = 4) were relatively simple (#1-3 and 5-8, *k* = 3) (Fig. 3A). The Heip power function (#2) was the most commonly-used function for Crustacea (9.4 % of all datasets, or 35.1 % of crustaceans), followed by the Bĕlehrádek (#8, 6.6 % of datasets or 24.6 % of crustaceans) and Tauti or exponential (#7, 4.7 % of datasets, 17.5 % of crustaceans) functions (Fig. 3A). The distribution of best functions as concluded in reviewed studies showed a similar pattern, with the linear rate function (#5) dominating for insects and arachnids and the Heip power function (#2) most often being concluded best for crustacean data (Fig. 3B). Among insects and arachnids, the Brière-1 function (#9) showed a slight tendency to be selected as best more often than other nonlinear functions, as it was for 7.0 % of all datasets, representing 6.7 % of insects and 18.2 % of arachnids (Fig. 3B).

### 3.2 Results of meta-analysis – did previous studies use the “best” function for their data?

The development function(s) used to fit species datasets and/or concluded to have been the “best” function for these data in their original studies were found, in the vast majority of cases, not to be the best function for the data out of the 33 functions examined (Fig. 4). When AIC_C_ was used to rank functions, the best function for 91.1 % of 79 retested study datasets was concluded to be a different one than that selected in previous studies (Fig. 4A), and for 86.1 % of datasets the actual best function was not even included in the set of all functions used in original studies (Fig. 4B). The actual best function was found to be different from that concluded to be best in original studies for all (100 %) arachnid and insect studies examined, and for most (75.0 %) crustacean studies (Fig. 4A). Also, for all taxonomic groups a considerable majority of datasets (100 % of Arachnida, 67.9 % of Crustacea, and 95.1 % of Insecta) were found to be best fit using a function that was not used in original studies of these species (Fig. 4B).

**Figure 4.**
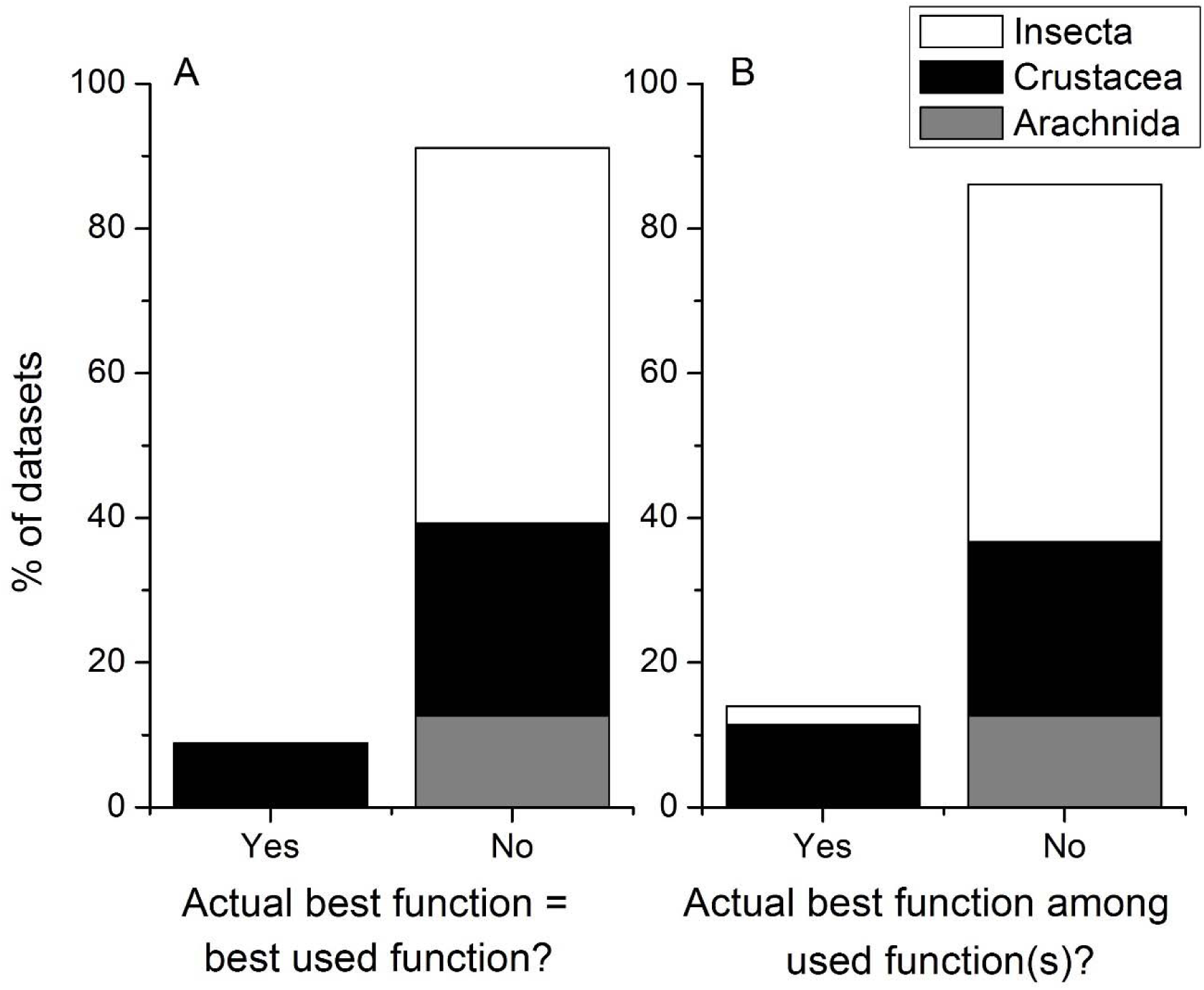
Percentage (%) of species datasets re-analyzed for meta-analysis (n = 79) for which the development function found to be best for the data (lowest AIC_C_ value, ranking based on AIC_C_ = 1) in this study (i.e., actual best function) (A) agreed or not with the best function as used in its original published study or (B) was among the set of all function(s) used within its original published study. Results are also broken down by taxonomic group, as in Fig. 2. For names and details of development functions (&1-33) the reader is referred to Table 1.

### 3.3 Overall performance of different functions

No one function was found to always be the “best” or “worst” for all reanalyzed datasets, but some functions did tend to perform better than others (Fig. 5). Those that were ranked as the best function by AIC_C_ particularly often (i.e., for ≥ 10 % of datasets) were the Heip power (#2), hyperbola (#3), Bĕlehrádek (#8), quadratic time (#16), and 3^rd^ order polynomial time (#19) functions (Fig. 5). There were no clear taxonomic patterns in terms of which functions tended to be ranked best, although function #19 did perform particularly well for insect datasets, and more complex functions with *k* > 5 were rarely concluded to be best (Fig. 5). Of the 33 functions tested, 16 were also never concluded to be the best of those tested (Fig. 5).

**Figure 5.**
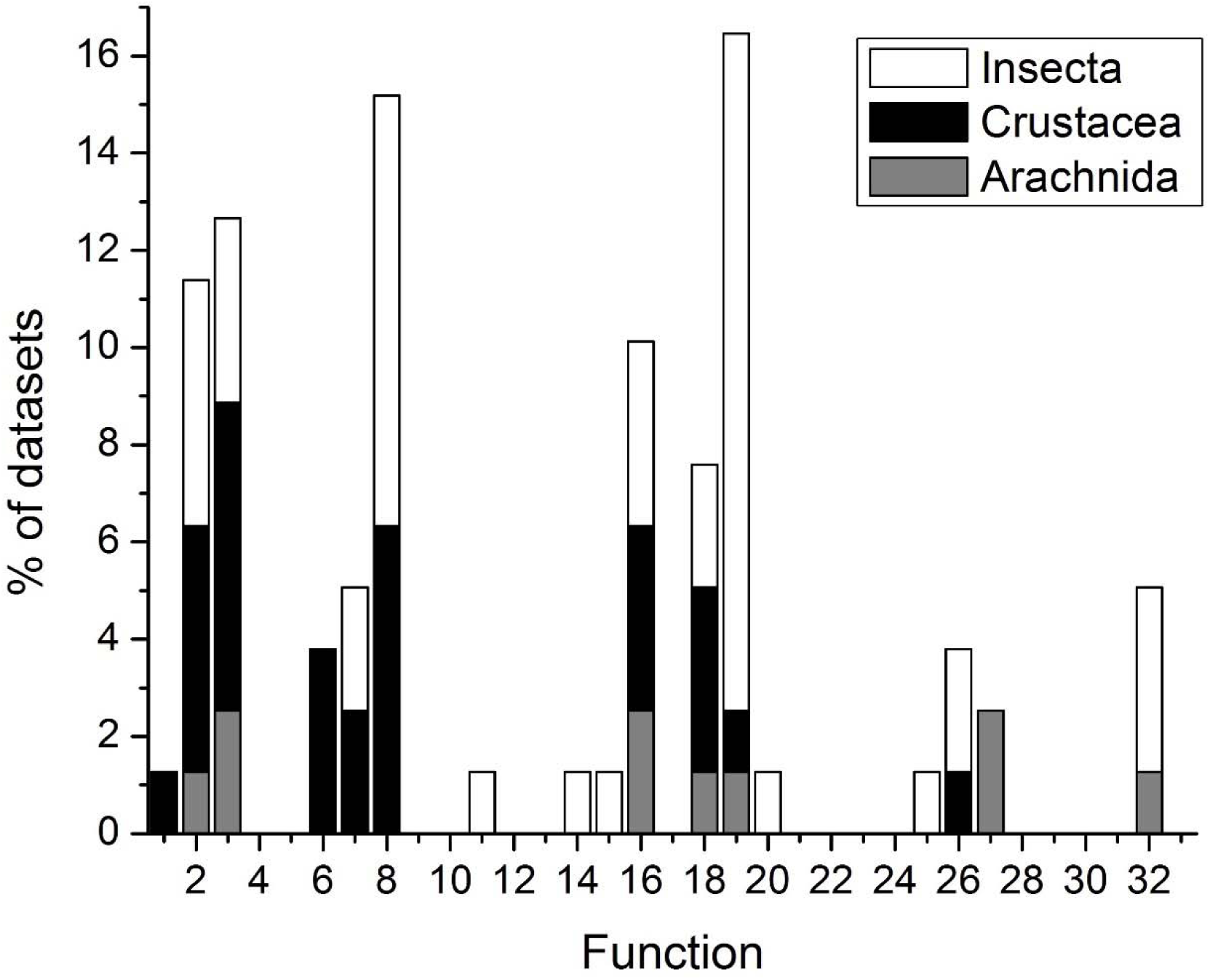
Percentage (%) of species datasets re-analyzed for meta-analysis (n = 79) for which each development function type was found to be best for the data (lowest AIC_C_ value, ranking based on AIC_C_ = 1) in this study. Results are broken down by taxonomic group, as in Fig. 2. For names and details of development functions (&1-33) the reader is referred to Table 1.

There were significant interactions between the effects of taxonomic group and development function type on overall performance of the 33 different development functions as assessed with R^2^-values (F _64, 2508_ = 1.377, p = 0.026) and by ranking functions with AIC_C_ (F _64,_ 2508 = 2.362, p < 0.001). Therefore differences in R^2^ and rank among functions were compared for each taxonomic group separately. In all three taxonomic groups, R^2^-values (Arachnida: F _32,_ 297 = 1.783, p = 0.007; Crustacea: F _32, 891_ = 3.721, p < 0.001; Insecta: F _32, 1320_ = 2.868, p < 0.001) and ranks significantly differed among different development functions (Arachnida: F _32, 297_ = 5.724, p < 0.001; Crustacea: F _32, 891_ = 14.707, p < 0.001; Insecta: F _32, 1320_ = 17.823, p < 0.001). However, overall Δ_i_ values were found to not differ significantly among functions (F _32, 2508_ = 0.278, p > 0.999) or taxonomic groups (F _2, 2508_ = 1.0, p = 0.368), nor was there a significant interaction between the effects of these factors on Δ_i_ values (F _64, 2508_ = 0.448, p > 0.999).

Differences in fit (R^2^) and rankings among functions were actually very similar across the different taxonomic groups (Fig. 6A-F; Table 2). Most function had relatively high overall average R^2^-values between 0.7 and 0.9 or higher (Fig. 6A-C), so on average, all temperature-dependent development functions tested were able to explain the majority of variation in observed development times. Functions with notably lower fit compared to others (lowest mean = 0.476) did occur, though, and included the linear time (#6) Logan-6 (function #25), Logan-10 (#30), and W-L-D (#33) functions on arachnid and insect data (Fig. 6A, C; Table 2), and the Ratkowsky (#27), Brière-1 (#9) and Brière-2 (#20) functions on crustacean data (Fig. 6B; Table 2). The explanatory power of development functions therefore differed by as much as ca. 10-50% on average depending on which was used (Fig. 6A-C). Functions with lower (better) overall ranks for all taxonomic groups included the Heip (#2), Hyperbola (#3), Bĕlehrádek (#8), quadratic time (#16), and 3^rd^ order polynomial time (#19) functions (Fig. 6D-E; Table 2), and those with high (poor) overall ranks included those with low R^2^-values described above and the Holling Type III (#21) functions (Fig. 6D-E; Table 2). All mean Δ_i_ values were high (mean Δ_i_ ≥ 22.7) because each function was not selected as the best function by AIC_C_ at least once, and in many cases the difference between the AIC_C_ values of the best function and the 2^nd^ best function (Δ_i_) were quite large (as evidenced by the variance in Fig. 6G-I). There were therefore large differences in the amount of information attained depending on which function was used, but no clear pattern among functions or taxa (Fig. 6G-I).

**Table 2.** Results of post-hoc comparisons of fit (R^2^) and rank (based on comparisons of AIC_C_ values) among different development functions within each arthropod subphylum using Tukey’s HSD test. Functions with different letters in a particular column had significantly different values of R^2^ or rank; comparisons were not made among subphyla (i.e., among columns). Functions labelled with the letter ‘A’ had the overall best performance (highest mean R^2^, rank closest to 1) while letters from B to M indicate progressively poorer function performance. Means of the R^2^ and rank data compared in this table are plotted in Fig. 6. Post-hoc comparisons were not made of Δ_i_ values because these did not differ significantly overall among functions. For names and details of development functions (&1-33) the reader is referred to Table 1.

| Function | $R^2$ | | | Rank | | |
| --- | --- | --- | --- | --- | --- | --- |
|  | Arachnida | Crustacea | Insecta | Arachnida | Crustacea | Insecta |
| 1 | AB | ABCD | ABCD | BCDEF | CDEFGH | EFGHIJ |
| 2 | A | AB | AB | ABC | A | ABCD |
| 3 | A | AB | AB | A | A | AB |
| 4 | AB | ABCD | ABCD | BCDEF | CDEFGH | FGHIJKL |
| 5 | AB | ABC | ABCD | ABCDEF | CDEFG | FGHIJKL |
| 6 | B | ABC | ABCD | DEF | DEFGH | LM |
| 7 | AB | ABC | ABC | ABCDE | ABC | DEFGH |
| 8 | A | AB | A | A | A | A |
| 9 | A | D | ABCD | ABCDEF | EFGHI | CDEF |
| 10 | AB | ABC | ABCD | ABCD | ABCD | BCDE |
| 11 | A | ABCD | ABCD | BCDEF | CDEFGH | DEFGHI |
| 12 | AB | ABCD | ABCD | ABCDEF | DEFGH | GHIJKLM |
| 13 | AB | ABC | ABCD | CDEF | CDEFGH | EFGHIJ |
| 14 | A | ABC | ABCD | ABCD | ABCD | BCDE |
| 15 | AB | ABC | ABCD | CDEF | ABCDE | EFGHIJKL |
| 16 | A | A | AB | AB | AB | ABC |
| 17 | A | ABC | ABCD | ABCDE | ABCD | BCDE |
| 18 | A | ABC | ABCD | ABCDE | BCDEF | EFGHIJ |
| 19 | A | ABC | ABCD | ABC | ABCD | BCDE |
| 20 | AB | CD | ABCD | ABCD | EFGHI | BCDE |
| 21 | A | ABCD | ABCD | EF | HI | JKLM |
| 22 | AB | ABC | ABC | ABCDEF | CDEFGH | EFGHIJKL |
| 23 | AB | ABC | ABCD | BCDEF | EFGHI | GHIJKLM |
| 24 | AB | ABCD | ABCD | ABCDEF | FGHI | EFGHIJKL |
| 25 | AB | ABCD | CD | BCDEF | EFGHI | KLM |
| 26 | A | AB | ABCD | ABC | ABCDE | BCDE |
| 27 | AB | BCD | ABCD | ABCDE | GHI | EFGHIJ |
| 28 | A | ABC | ABCD | ABCDEF | CDEFGH | EFGHIJK |
| 29 | AB | ABC | ABC | ABCDEF | CDEFGH | DEFG |
| 30 | B | ABC | BCD | F | HI | M |
| 31 | AB | ABC | ABCD | BCDEF | GHI | HIJKLM |
| 32 | AB | ABC | ABC | ABCDE | EFGHI | DEFG |
| 33 | AB | ABCD | D | EF | I | IJKLM |

**Figure 6.**
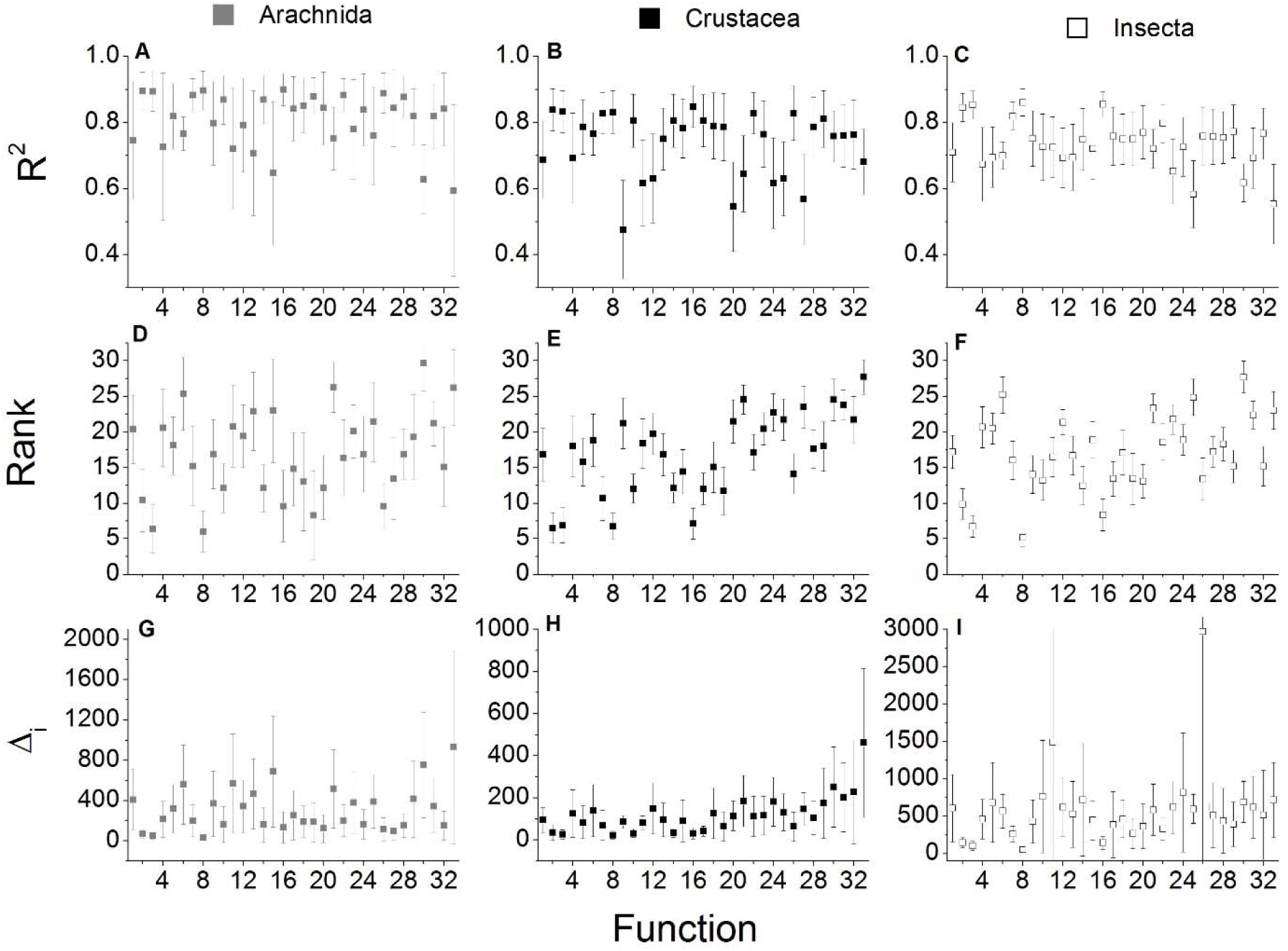
Overall performance of different development functions (x-axes) assessed based on (A-C) R^2^ values, (D-F) ranking based on AIC_C_ values, and (G-I) Δ_i_ values (y-values). Function performance was assessed separately for each taxon: (A, D, G) arachnids (n = 10 species for each function), (B, E, H) crustaceans (n = 28), and (C, F, I) insects (n = 41). Possible rankings ranged from 1 (“best” model) to 33 (“worst”). Values plotted are mean values per function and taxonomic group taken across all species datasets within that group ± 95 % C.I.s. For names and details of development functions (&1-33) the reader is referred to Table 1. Results of post-hoc comparisons among functions with Tukey’s HSD test for each taxonomic group are presented in Table 2.

### 3.4 Consequences of function choice

All measures of prediction error, decreased fit, and increased information loss resulting from using the best original rather than the actual best functions on species datasets were significantly different from zero (one-sample *t*-tests, p < 0.05; Table 2). Development times predicted with original studies’ functions disagreed with observed developmental duration by about 4 days or 85.9 % on-average, but could be off by as much as 132 days or 1725.5 % (Table 3). Fit (R^2^) of original functions to arthropod datasets was lower by 0.091 on-average compared with best functions (Table 3), meaning that nearly 10 % of the variation in the data would remain unexplained if the original rather than the actual best function was used. The difference in R^2^-values between originally-used and best functions could be much greater in many cases, though, as percent decrease in R^2^ by using the best function was as high as 100 % for certain study datasets (Table 3; Table S2). Additionally the mean Δ_i_ of the originally-used functions was 225 and could be as high as 3958.7 for some datasets (Table 3), indicating a substantial loss of information relative to the actual best function as determined in the present study.

**Table 3.**
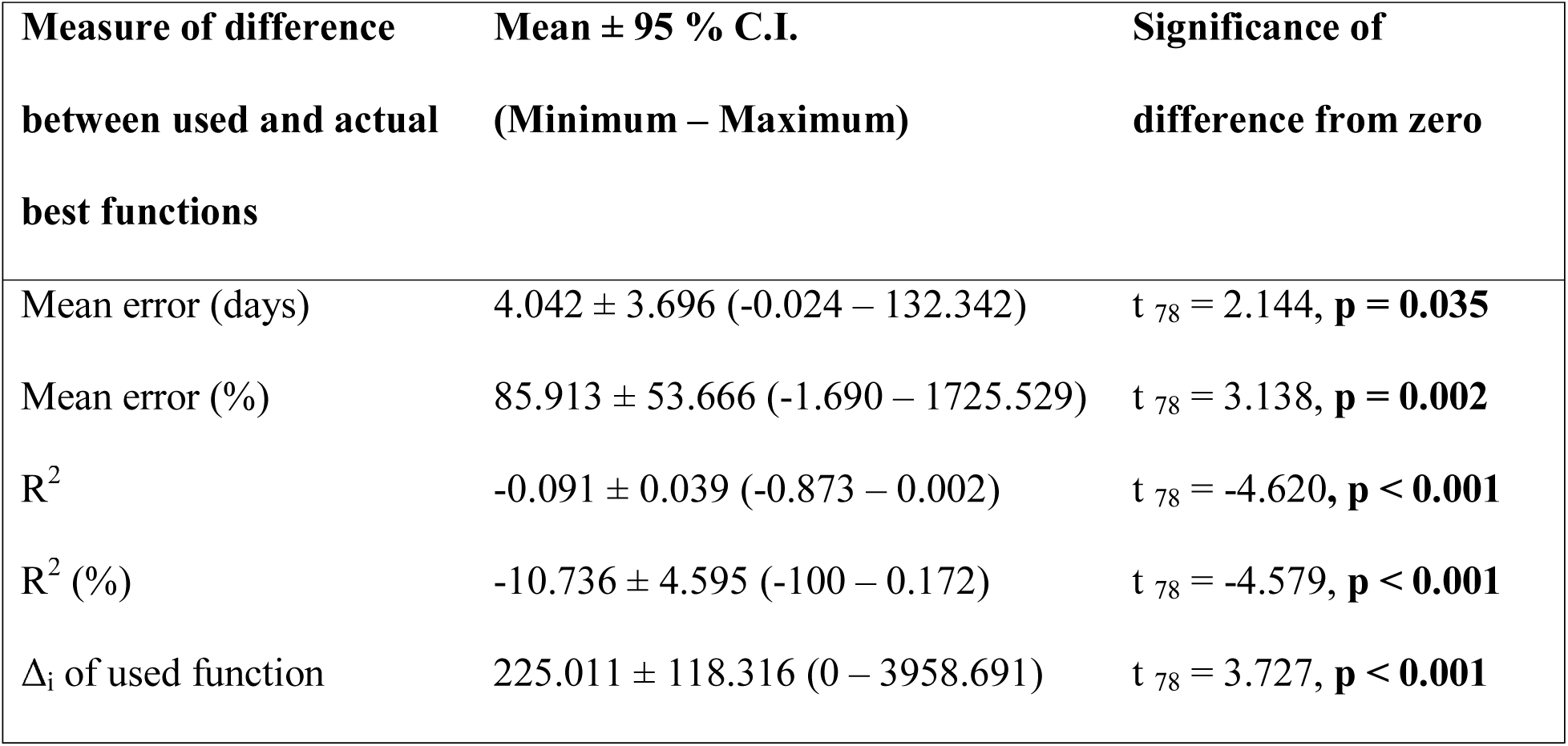
Consequences of fitting development data with functions used or found to be best in original published studies rather than the best function for each of 79 reanalyzed species datasets, as determined in this study. Consequences were assessed by calculating differences between used and actual best functions in terms of (1) increased mean error (absolute, in days) between observed and predicted development times, (2) increased percent (%) prediction error, (3) usually poorer fit (R^2^), (4) poorer percent (%) fit (R^2^), and (5) information loss (Δ_i_ of used models; Δ_i_ = 0 is the best model). resulting. Mean values ± 95 % confidence intervals (95 % C.I.) for each difference measure taken across all retested datasets are shown (with the range of values in parentheses), as well as the results of one-sample t-tests comparing these differences to zero; statistically-significant p-values (p ≤ 0.05) are presented in bold text.

| **Measure of difference**<br>**between used and actual**<br>**best functions** | **Mean ± 95 % C.I.**<br>**(Minimum – Maximum)** | **Significance of**<br>**difference from zero** |
| --- | --- | --- |
| Mean error (days) | 4.042 ± 3.696 (-0.024 – 132.342) | $t_{78} = 2.144$, **p = 0.035** |
| Mean error (%) | 85.913 ± 53.666 (-1.690 – 1725.529) | $t_{78} = 3.138$, **p = 0.002** |
| $R^2$ | -0.091 ± 0.039 (-0.873 – 0.002) | $t_{78} = -4.620$, **p < 0.001** |
| $R^2$ (%) | -10.736 ± 4.595 (-100 – 0.172) | $t_{78} = -4.579$, **p < 0.001** |
| $\Delta_i$ of used function | 225.011 ± 118.316 (0 – 3958.691) | $t_{78} = 3.727$, **p < 0.001** |

### 3.5 Range of temperature tested in studies versus function performance

Thermal ranges tested in original studies differed significantly among studies of different taxonomic groups (F _2, 96_ = 16.533, p < 0.001; see also Table S2). Interestingly, studies of crustaceans tended to be carried out over significantly smaller thermal ranges (mean ± 95% C.I. = 11.9 ± 0.8 °C, range = 6.0-22.5°C) than those of arachnids (20.0 ± 1.1 °C, range = 12.5-30.0°C; Tukey’s HSD test, p < 0.001) or insects (17.5 ± 1.2 °C, range = 8.0-38.0; Tukey’s HSD test, p < 0.001) implying notable differences in the way thermal effects are investigated in these taxa; thermal ranges studied for Insects and Arachnids did not differ, however (Tukey’s HSD test, p = 0.328).

The range of temperatures examined in previous studies was significantly correlated with performance (ranking) of 13 of the 33 development functions tested in this study (Fig. 7). With one exception (function # 30), all functions for which rankings by AIC_C_ were significantly and positively correlated with temperature range were those (#1, 2, 5-7, 13, 15, and 16) that had fewer parameters (*k* ≤ 4) (Fig. 7). This means that these relatively simpler functions tended to do more poorly (higher value = poorer rank, further from 1) when larger thermal ranges were tested. Conversely, rankings of more complex functions with *k* = 5 (#20, 24, and 27) or *k* = 7 (#32) were negatively correlated with temperature range (Fig. 7), meaning that these functions performed better (lower value = better rank, closer to 1) when studies tested a wider range of temperatures.

**Figure 7.**
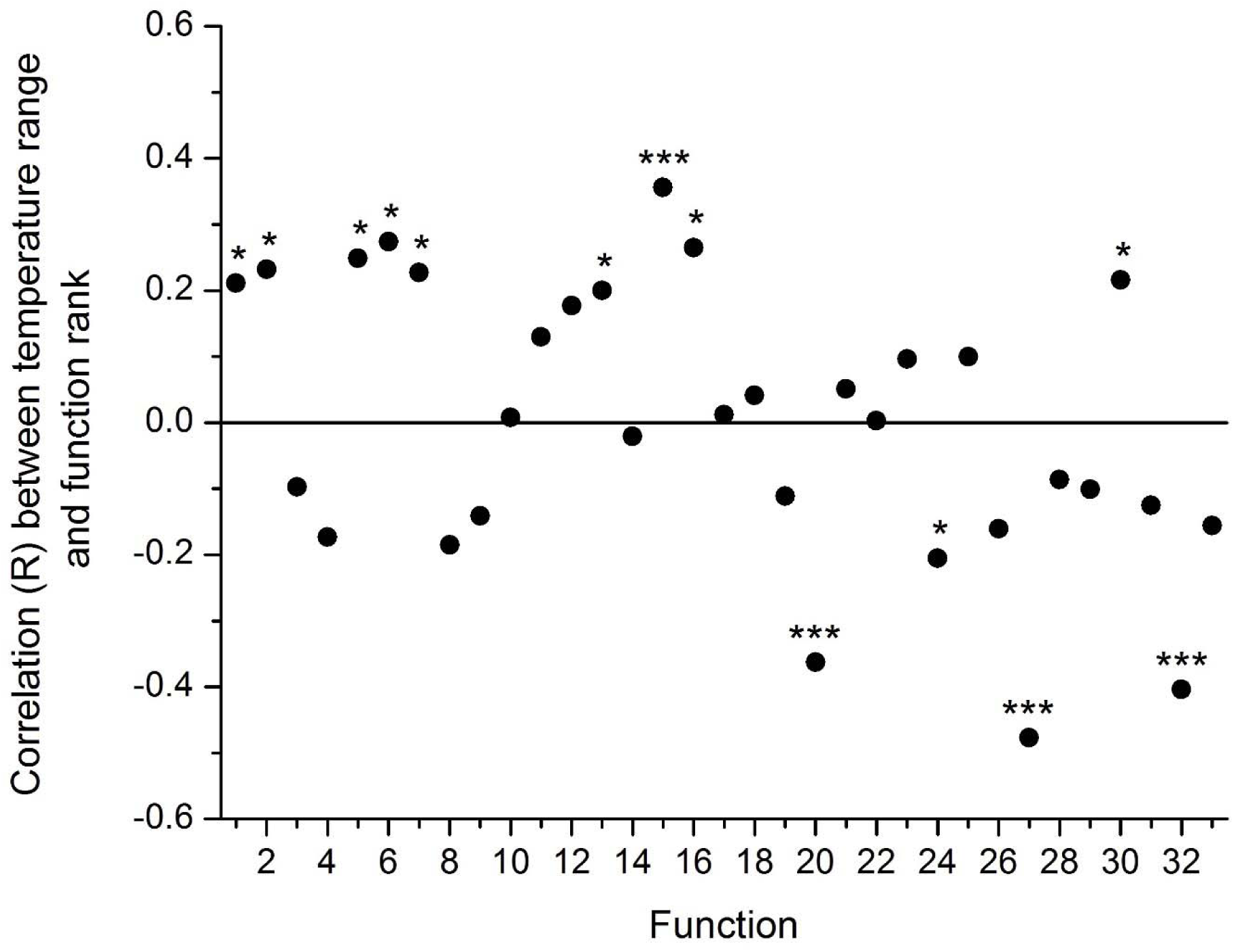
Pearson’s correlation coefficient values (R) calculated between the ranges of temperatures (°C) tested in all original studies (n = 99) and rank (out of 33) of development functions based on AIC_C_ for each species dataset. Statistical significance of correlation coefficients is indicated by labels above each plotted point as follows: * p ≤ 0.05, ** p ≤ 0.01,*** p < 0.001; non-significant (p > 0.05) results are not labelled. For names and details of development functions (&1-33) the reader is referred to Table 1.

## 4. Discussion

### 4.1 Use and performance of different development functions in previous studies

Modeling functional relationships between temperature and life history characters is an essential component of studying the biology of poikilothermic organisms (Angilletta Jr., 2006; Bĕlehrádek, 1935; Papnikolaou et al., 2013; Shi and Ge, 2010). Predictions of generation times (Huntley and Lopez,1992), timing of seasonal events (Bayoh and Lindsay, 2003), dispersal potential (de Rivera et al., 2007), and recruitment to adult populations of Arthropoda (Aiken and Waddy, 1986; Caddy, 1986) produced by such modeling efforts are thus sensitive to the types of temperature-dependent larval development functions incorporated in these. Development times of arthropod larvae predicted for the same species and temperatures by different function types can differ substantially, which has important impacts on predictions made. Using the best possible function to represent a given species’ and/or study’s dataset should thus be a crucial component of the study of temperature-dependent arthropod larval development, which should precede reporting and use of the results of such studies in models. However, in the present study this important step was found to be largely bypassed by the majority of studies. Particular function types tended to be used more often than others for particular taxonomic groups with little or no clear justification for the choice made, while consideration of alternative function types was rarely reported in published papers. In most cases the function used in original published studies was not actually the best one for the datasets presented. Further, fitting these data with the best model resulted in better fit, less disagreement between predicted and observed development times, decreased information loss, and presumably also better predictive ability. These results demonstrate that development function choice is an important but often-ignored step in research on arthropod larval development, which should be given greater consideration in future studies.

Choosing one particular development function might have some justification if any function(s) could be said to be better overall than others. In the present study, no single function type was found to be the best or worst, although some did tend to perform better or worse than others (see Fig. 5, 6 in Results). Functions that performed well overall might be recommended as good starting points for fitting development data, and those that did poorly overall could conversely be used with caution. (Table S1). Also, the more complex functions with high *k*-values and including T_min_ and T_max_ parameters, which performed poorly overall, did somewhat better on insect data than for other taxa and actually was among the best models for some insect species datasets (Table S2). Therefore, it is difficult to make general statements about which function is always best to use; this must rather be assessed on a case-by-case basis, for each species and study. Results in this study showed that not using the best function for a given dataset can result in very different predicted development times, which could lead to very different (and potentially erroneous) inferences and predictions of species biology by modeling studies (Miller et al., 1998; Miller et al., 2006; Quinn, 2014; Reitzel et al., 2004). The practice among many fields of study has been to fit data with a development function type that has been used in previous studies on similar species; for example, the frequent use of the Bĕlehrádek function on copepod crustaceans (Anger, 2001; Corkett and McLaren, 1970; Hamasaki et al. 2009) or linear rate + complex function(s) on insects and arachnids (Golizadeh and Zalucki, 2012; Shi and Ge, 2010; Smits et al., 2003; Table S2). However, based on results of the present study this practice should be discontinued.

### 4.2 Importance of temperature range tested to function performance

An interesting finding in the present study was that overall performance of several function types was correlated with the range of temperatures tested in original published studies. Specifically, as the range of temperatures tested increased performance (i.e., likelihood to be ranked as the best model) of the simpler functions examined decreased while that of more complex functions increased. This result does make sense, however, if one considers the “real” nature of temperature-biological rate relationships. Because the actual performance of the enzymes mediating larval development most certainly have upper and lower functional threshold temperatures, beyond which development cannot progress (Brière et al., 1999; Quinn and Rochette, 2015; Somero, 2004), one can assume that for most species the “true” relationship between temperature and development time resembles the Brière-2 function, or similar complex asymmetrical curves (e.g., Huey and Stevenson, 1979; Shi and Ge, 2010). A study carried out over a very wide range of temperatures should be able to approach or exceed thermal thresholds and therefore identify these limiting temperatures, and thus be best explained by a complex function. However, if one carries out their study over a more narrow thermal range, they will only be able to observe a certain section of the development curve, which could be located relatively far from one or both threshold temperatures. This could result in the observed temperature-development data having a distinctly linear, quadratic, or power function-like shape, such that one of these alternative, simpler functions would be identified as the “best” function for the data over this specific range. Indeed, this seems to have been the case in several of the datasets examined, in which thermal ranges and/or sample sizes were relatively small (e.g., Quinn et al., 2013; Corkett and McLaren, 1970; Hamasaki et al., 2009; Carlotti et al., 2007; Table S1, S2) and the best function was determined to be one of the simpler functions, such as the linear time (#6) or Heip power (#2) function, even though these should be the least-realistic (Bĕlehrádek, 1935; Brière et al., 1999; Somero, 2004). Importantly, when this occurs the best function will be the one that provides the most informative description of development times over a very specific range of temperatures, but its performance is likely to degrade if extrapolation beyond this range is attempted. Ultimately the “best” function should be of a complex form resembling functions with T_min_ and T*max* parameters, but most studies, especially of Crustacea, are not conducted over sufficiently wide temperature ranges to be allow good estimates of such functions’ parameters to be derived.

The vast majority of insects and arachnids have terrestrial and/or freshwater aquatic habitats, in which temporal variability in air and water temperatures can be very large (Pakyari et al., 2011; Sanchez-Ramos et al., 2007; Stavrinides et al. 2010). As a result, the likelihood of these organisms and their larvae being exposed to extreme temperatures exceeding thresholds for moulting, development, and/or survival can be high. Conversely, many crustaceans (and all of those examined in the present review; Table S2) inhabit the marine realm as larvae and/or adults (Paul and Paul, 1999; Roberts et al., 2012; Thompson, 1982). While it is not impossible that marine crustacean larvae could encounter temperatures too high for development or survival (e.g., such warm extremes could occur in shallow coastal areas, highly-stratified water columns, intertidal zones at low tide, or more generally due to future climate change; Caffara et al., 2012; Quinn and Rochette, 2015), they are thought to be far more likely to encounter lower limiting temperatures, especially in the deeper ocean or temperate regions (Hartnoll, 1982; MacKenzie, 1988; Quinn, 2016). This perceived difference in limiting temperatures appears to have lead studies on temperature-dependent development in these groups along different paths. Studies of insects and arachnids often use very wide temperature ranges with the intent of capturing lower and upper limiting temperatures for development in their species because these physiological limits are known to be essential to modeling these species in their natural environments (Shi and Ge, 2010; Smits et al., 2003). Studies of crustaceans, to the contrary, tend to be limited to more narrow thermal ranges deemed “ecologically-relevant” (i.e., likely to be encountered by the species in nature); occasionally these include lower limits, but in general physiological limits, especially upper ones, are rarely sought (Hartnoll, 1982; Quinn, 2016).

While there is certainly logic behind the use of narrower, more-relevant thermal ranges in studies of Crustacea, this approach still has potential to result in errors for two main, related reasons. First, the type of development function used changes predicted development times both at and between (interpolation) observed temperatures and especially outside of these (extrapolation) (Angilletta Jr., 2006; Campbell et al., 1974; Quinn and Rochette, 2015). Second, thermal development limits actually change the shape of the “real” and estimated (i.e., fitted by regression) development curve, for example by decreasing its curvature when lower and upper limits are further apart and increasing curvature when these are closer together (Bĕlehrádek, 1935; Brière et al., 1999; Shi and Ge, 2010; personal observations by author). All else being equal, these difference in curvature can result in very different development times at the same temperatures. As a result, it is important to know the physiological limits of a given species when modeling its development (Quinn, 2016). Even if a species rarely encounters temperatures close to these limits, development times calculated at intermediate temperatures will be impacted by the values of these limits; if one ignores these limits and uses a different function type, or attempts to estimate limits by extrapolation from a narrow thermal range, there is great potential for erroneous predictions of development times to be made.

### 4.3 Discussion of potential limitations and next steps

In the present review, published studies on arthropod larvae were obtained through literature searches through Web of Science (Thompson Reuters, 2015). These searches were by no means comprehensive – many other studies of temperature-dependent development of arthropod larvae exist that were not indexed in this search engine – but it was extensive and did provide a good sample of such studies encompassing many different years, regions, and arthropod taxa (Table S2). This sample of the relevant literature was thus appropriate and useful for the purposes of the present review of development function usage and performance. An expanded search using additional search tools in a future study could obtained data for other taxonomic groups within the Arthropoda (e.g, myriapods, more arachnids, other orders of Crustacea and Insecta, etc.); indeed, an Insect Developmental Database has been created by Nietschke et al. (2007) that could be used to obtain considerably more insect data for reanalyses. Performance of development functions on data from species outside of the arthropod phylum (e.g., molluscs: de Severyn et al., 2000; nematodes: Jenkins et al., 2006; Singh and Sharma, 1994; urochorates: Kang et al., 2009; vertebrates: McLaren and Cooley, 1972; Miller et al. 2006) may also be attempted, and reveal additional patterns in study design, taxonomy, and function usage of interest. However, overall patterns and conclusions of the present study would likely hold true. To simplify analyses, this study conducted analyses on total larval development time data rather than on individual larval stages. However, in most species development time of each larval stages has a distinct response to temperature, requiring different developmental equations to be derived for each stage (Corkett and McLaren, 1970; Hartnoll, 1982). Often survival to and through later larval stages is very low, so power to fit more complex functions to later-stage development data can be limited. As a result, the best function can potentially differ among larval stages of the same species, in the same study; indeed, this was noted for American lobster data in the present study (data not shown). A future study should investigate stage-specific changes in the “best” development function(s), to confirm whether such patterns could impact the overall performance of different function types, prediction error, and so on. However, it would make mathematical sense for similar patterns to be found through such a detailed review to those noted in the present study, given that similar factors (e.g., sample sizes and thermal ranges tested) would impact function performance. There are also countless other development function types in existence which were not included in the present study (e.g., Angilletta Jr., 2006; Schoolfied et al., 1981; Shi and Ge, 2010). It is possible that one or more of the functions not examined herein could actually be the closest to “real” development relationships and/or perform better overall than all others. However, findings of this review that one or few functions are used by most studies, the best function was not the one used in most original published studies, and that using the non-best function results in poorer predictions would not change through consideration of such additional functions.

In this study function performance was assessed mainly in terms of fit (R^2^ and observed versus predicted values) and information loss (AIC_C_ and Δ_i_). These gave good indications of how appropriate each function and its parameter estimates were for particular datasets (e.g., how well sample sizes supported estimation of more complex functions). However, future studies could take more thorough approaches to assessing predictive ability of different functions. One approach to be used in the future to assess function performance could be cross-validation (Picard and Cook, 1984; Anderson, 2008). Even better would be actually testing development functions on new data, for example by predicting development times for a particular species at different temperatures using different functions derived in a prior study, and then measuring new development times and comparing these to predictions. If future studies attempted this, very thorough tests and new evidence in favour of one function or another might be obtained.

### 4.4 Implications of development time predictions based on different functions

Differences in development times predicted for the same species and temperature among development functions have potential to impact various types of predictions relevant to arthropod biology and ecology. Larval survival is usually inversely related to larval duration, such that slower development results in fewer potential recruits to adult populations (Reitzel et al., 2004; Roberts et al., 2012). Most life history and bio-physical models account for this by reducing larval numbers in simulated cohorts by a certain percentage at each model time step, resulting in substantial, exponential losses per each additional step spend in larval development (e.g., Miller et al., 1998; Quinn, 2014). In many crustaceans, the larval phase of the life cycle is the main dispersive phase (Anger, 1984; de Rivera et al., 2007; MacKenzie, 1988). Lengthening the larval developmental period of such larvae can dramatically alter how far and to where larvae drift in simulations with ocean currents; for example, simulations by Quinn (2014) showed that slowing larval development (and thus lengthening drift time) of American lobster larvae by 60 % could result in increased drift distances of larvae by up to ca. 500 km. If drift were overestimated in such models, for instance due to use of inappropriate development functions, then the degree of population mixing would be overestimated and an incorrect estimate of population structure obtained; underestimation by the same means could also lead to considerable errors. Likewise if dispersal ability of larvae of an invasive species, such as the green crab *Carcinus maenas* (L.) (de Rivera et al., 2007), were underestimated in this way potential invasions to new regions that could be predicted may be missed. Using development functions to estimate the timing of seasonal peaks in abundance of disease vectors (Bayoh and Lindsay, 2003), agricultural pests (Campbell et al., 1994; Easterbrook et al., 2003; Stavrinides et al., 2010), or species that serve as important food sources to others (e.g., copepod secondary productivity in the ocean: Carlotti et al., 2007) also depends on being able to make good estimates of larval development. For many species very small differences in development time similar to the difference in “errors” of best and original studies’ functions are enough to dramatically alter the nature and implications of modeled predictions (e.g., ≤ 1-5 days; Gadino and Walton, 2012). The type of development function used can thus have large impacts on predictions, so it is important that studies attempt to find the best model for their data. Importantly, much research is now being initiated to assess how future climate change will impact many species, including arthropods and their larvae (Caffara et al., 2012; Quinn and Rochette, 2015). Use of non-best development functions within such research clearly could result in erroneous predictions as well and so should be avoided.

### 4.5 Recommendations and conclusions

Based on results of this study, it is recommended that future studies examining effects of temperature on development of arthropod larvae consider and attempt to fit multiple alternative development function types to their data to determine the best way to model their results and report that this was attempted. No one function type is better or worse overall, but the range of temperatures to be considered and potential use of results (e.g., for extrapolation to temperatures not observed) can be used as a guide when deciding which functions are most likely to provide good representations of data. In general simpler functions could provide better descriptions of development observed over relatively narrow thermal ranges, but provide poor extrapolation ability. Conversely, wider ranges encompassing lower and/or upper limiting temperatures for development can support more complex functions, which potentially resemble more closely true enzymatic and biological thermal performance curves (Brière et al., 1999; Somero, 2004) and may allow modeling over all temperatures potentially encountered. Studies on crustaceans in particular should be conducted over wider thermal ranges in the future so that limiting temperatures of these species can be identified and more complex, presumably realistic development functions reliably fit to data for these organisms. Considering different potential function types to find the best for each dataset should lead to better predictions of larval development times in support of subsequent research on these important species.

## Acknowledgements

Thanks are due to Rémy Rochette, Joël Chassé, Jeff Houlahan, and Heather Hunt for advice and guidance during this project, and the University of New Brunswick, Saint John Campus, for providing resources that made the review and analyses possible. The author also thanks two anonymous reviewers for providing comments that improved the scope and quality of the manuscript.

## Electronic Supplementary Material

**Table S1.** Development function parameter estimates, R^2^, and AIC_C_ for American lobster data plotted in Figure 1.

**Table S2.** List of studies and species obtained for the literature review and meta-analyses, including taxonomic grouping, function type(s) used, best function, whether the best and used function were the same, temperature range tested, performance (R^2^, AIC_C_ rank, and Δ_i_) of each of the 33 functions tested (‘na’ = study not possible to retested in meta-analysis), and consequences of using original studies’ rather than the actual best function to fit datasets in terms of prediction error, R^2^ decrease, and information loss (Δ_i_).

